# Planning and navigation as active inference

**DOI:** 10.1101/230599

**Authors:** Raphael Kaplan, Karl J Friston

## Abstract

This paper introduces an active inference formulation of planning and navigation. It illustrates how the exploitation–exploration dilemma is dissolved by acting to minimise uncertainty (i.e., expected surprise or free energy). We use simulations of a maze problem to illustrate how agents can solve quite complicated problems using context sensitive prior preferences to form subgoals. Our focus is on how epistemic behaviour – driven by novelty and the imperative to reduce uncertainty about the world – contextualises pragmatic or goal-directed behaviour. Using simulations, we illustrate the underlying process theory with synthetic behavioural and electrophysiological responses during exploration of a maze and subsequent navigation to a target location. An interesting phenomenon that emerged from the simulations was a putative distinction between ‘place cells’ – that fire *when* a subgoal is reached – and ‘path cells’ – that fire *until* a subgoal is reached.

## Introduction

The ability to navigate an uncertain world is clearly a central aspect of most behaviour. This ability rests on the optimal integration of knowledge about the world and the goals that we are currently pursuing (Hauskrecht, 2000, Johnson et al., 2007, Pastalkova et al., 2008, Hassabis and Maguire, 2009, Humphries and Prescott, 2010, Karaman and Frazzoli, 2011, Buzsaki and Moser, 2013, Pfeiffer and Foster, 2013a). This paper offers both a normative and process theory for planning and navigating in novel environments – using simulations of subjects performing a maze task. Our objective was not to find an optimal solution to the problem at hand; rather, to develop a model of how the problem could be solved in a neurobiologically plausible fashion. In other words, we wanted to establish a modelling framework within which we can compare different models in terms of their ability to explain empirical responses; i.e., reaction times, saccadic eye movements and neurophysiological responses. To accomplish this, we focus on a minimal model of nontrivial planning that involves navigating a maze from a start location to a target location. Crucially, we consider this problem under uncertainty about the maze – thereby requiring the subject to explore the maze (visually) and then use this information to navigate to the target or goal. In what follows, we describe an active inference scheme based on Markov decision processes that accomplishes this task. This paper restricts itself to describing the scheme and generative model – and to illustrating the model predictions, using simulated behavioural and electrophysiological responses. Subsequent work will use the model described in this paper to characterise empirical responses and compare different models of behaviour along the lines described in (Schwartenbeck and Friston, 2016).

The contribution of this work is not so much the solution to the maze problem but the sorts of solutions that emerge under Bayes optimality principles (i.e., active inference) when plausible constraints are applied: see also Solway & Botvinick (2015). These constraints range from general principles to specific constraints that must be respected by real agents or sentient creatures. For example, at a general level, we suppose that all perception (i.e., state estimation) and consequent behaviour conforms to *approximate* Bayesian inference – as opposed to *exact* Bayesian inference. In other words, by using a variational (free energy) bound on model evidence, we implicitly assume a form of bounded rationality. At a more specific level, realistic constraints on inference arise from how the environment is sampled and evidence is accumulated. For example, we will use synthetic subjects that have a limited working memory that can only entertain short-term (finite horizon) policies. Furthermore, we will use agents who have a rather myopic sampling of the environment, obliging them to forage for information to build a clear picture of the problem with which they are contending. These constraints, particularly the limited horizon of prospective planning, lead to, or mandate, a simple form of hierarchical planning. This basically involves identifying proximal subgoals – within reach of a finite horizon policy – that necessarily lead to distal goals in the long term; c.f., (Sutton et al., 1999).

This paper comprises three sections. The first section reviews active inference and the form of the generative models necessary to specify normative (uncertainty resolving) behaviour. It deals briefly with the underlying process theory, in terms of evoked electrophysiological responses and the associative plasticity of neuronal connections. The second section describes a particular generative model apt for solving the maze problem. This problem can be regarded as a metaphor for any sequence of constrained state transitions that have to be selected under uncertainty about the constraints. The final section provides some illustrative simulations to show the sorts of *in silico* experiments that can be performed with these sorts of schemes. This section concludes by considering how trajectories or paths through (state) spaces might be encoded in the brain. In particular, we consider the notion of ‘place cells’ and ask whether place-cell-like activity may be a subset of more generic ‘paths cells’ that report “where I have come” from, as opposed to “where I am”. We conclude with a discussion of how the active inference scheme described in this paper relates to – and inherits from – previous work in reinforcement learning and theoretical neurobiology.

## Active inference and resolving uncertainty

Over the past years, we have described active inference for Markov decision processes in a wide range of settings. These cover simple (two-step) choice tasks to complicated hierarchical inference; for example, in reading sentences (Friston et al., 2015, Mirza et al., 2016). The underlying principles of active inference do not change. The only thing that changes is the generative model that specifies the task or scenario at hand. What follows is a formulation of planning and navigation using the same scheme used previously to explain other perceptual, behavioural and cognitive phenomena.

The specific aspect of the current application rests upon how prior beliefs are specified. By showing that planning and navigation can be modelled with a generic (active inference) scheme, we hoped to show (i) that many aspects of planning transcend the particular problem of spatial navigation and (ii) the solutions that emerge speak to – and contextualise – previous formulations: e.g., (Sun et al., 2011a, Solway et al., 2014, Fonollosa et al., 2015, Maisto et al., 2015, Donnarumma et al., 2016, Stachenfeld et al., 2017, Gershman and Daw, 2017).

Active inference refers to the minimisation of surprise – or resolution of uncertainty – during the active sampling of an environment. Formally, active inference is a normative theory, in the sense that there is a single objective function; namely, variational *free energy.* Free energy provides an upper bound on *surprise* (the improbability of some sensory samples), such that minimising free energy implicitly minimises surprise. An alternative perspective on this optimisation follows from the fact that surprise is negative *model evidence*. In other words, active inference implies some form of self-evidencing (Hohwy, 2016); in the sense that inference and subsequent behaviour increase the evidence for an agent’s model of its world. Clearly, to make inferences, an agent has to entertain beliefs. In active inference, these beliefs are constituted by an approximate posterior density; namely, a probability distribution over the causes of sensory samples based on sensory evidence. The causes of sensory consequences are generally referred to as *hidden states* because they are generally hidden from direct observation and have to be inferred. In other words, active inference entails state estimation. Crucially, agents have beliefs about hidden states of the world *and their behaviour*. This sets active inference apart from other schemes, in the sense that inferences about action and behaviour become an integral part of the general inference problem. This enables state estimation and planning as inference (Attias, 2003, Botvinick and Toussaint, 2012) to be subsumed gracefully under a single objective; namely, self-evidencing.

Practically, actions are selected from posterior beliefs about sequences of actions or *policies*. Each action solicits a new observation from the world – leading to the next cycle of active inference or perception. There are two key aspects that fall out of this formulation. First, entertaining beliefs about sequences of action necessarily requires an agent to have beliefs about (i.e., approximate posteriors over) hidden states in the future (and past). This necessarily endows agents with a short term memory of the proximal future (and past) that can be used for prediction (and postdiction). The second key aspect is that *posterior* beliefs about policies rest on *prior* beliefs about future outcomes. These prior beliefs can be regarded as preferred outcomes or goals in a reinforcement learning or utilitarian (economics) setting. In short, the heavy lifting in active inference rests upon how prior beliefs about behaviour or policies are formed.

### Prior preferences, novelty and salience

In active inference, prior beliefs about policies are proportional to (negative) *expected* free energy. This follows naturally from the imperative to minimise surprise as follows: expected surprise is uncertainty (mathematically speaking, expected self-information is entropy). It therefore follows that surprise minimising (self-evidencing) policies must minimise expected surprise or, in bounded or approximate inference, they must minimise expected free energy. This is formally equivalent to choosing policies that resolve uncertainty (see Appendix 1 for a more technical description). When expected free energy is unpacked, several familiar terms emerge (Friston et al., 2015). The expected free energy for a particular policy at a particular time in the future can be expressed as (see Table 1 and Appendix 2 for a list of variables and technical description):

**Table 1:**
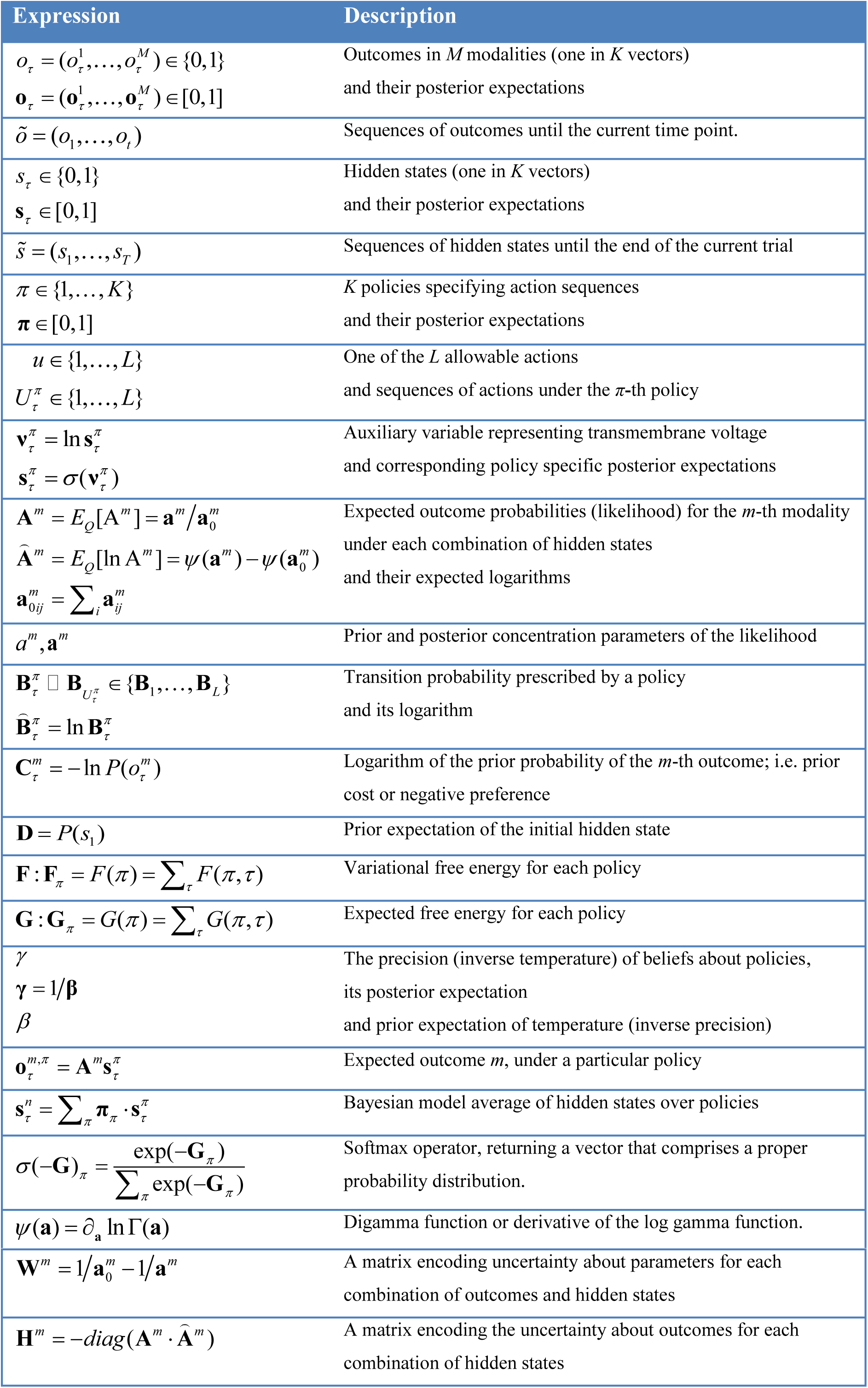
Glossary of variables and expressions

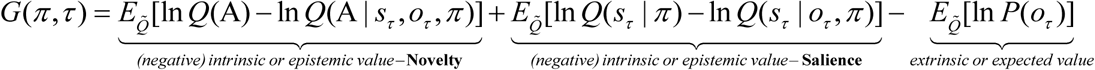

Here, 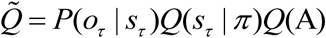 is the posterior (predictive) distribution over the probabilistic mapping A from hidden states *s_τ_* to outcomes *O_τ_* under a particular policy *π* at time *τ* in the future.

Intuitively, expected free energy can be divided into epistemic, information seeking and pragmatic, goal seeking parts, corresponding to *intrinsic* and *extrinsic* value respectively. Extrinsic (pragmatic) value is simply the expected value of a policy defined in terms of outcomes that are preferred *a priori*; where the equivalent cost corresponds to prior surprise. The more interesting parts are uncertainty resolving or epistemic in nature. These correspond to the first two (novelty and salience) terms above. These quantities are variously referred to as relative entropy, mutual information, information gain, Bayesian surprise or value of information expected under a particular policy (Barlow, 1961, Howard, 1966, Optican and Richmond, 1987, Linsker, 1990, Itti and Baldi, 2009). In short, they score the reduction of uncertainty that would accrue under a particular policy for sampling the world. In other words, they score the epistemic value of the evidence that would be accumulated by pursuing a particular sequence of actions.

Crucially, this uncertainty reduction comes in two flavours. There is an epistemic value associated with beliefs about the current state of the world – and how they unfold in the future. This epistemic value is generally referred to as the *salience* of sampling the world in a particular way: c.f., (Berridge and Robinson, 1998, Itti and Baldi, 2009, Friston et al., 2015). The equivalent salience for the parameters of a model (denoted by A) reflects the resolution of uncertainty about probabilistic contingencies that endows the world with causal structure. In other words, the epistemic value of a policy – that rests on uncertainty about model parameters (as opposed to hidden states) – encodes the *novelty* of a policy. Put simply, a novel situation becomes attractive because it affords the opportunity to resolve uncertainty about what would happen “if I did that”. In what follows, we will call upon novelty (the epistemic value of reducing uncertainty about model parameters) and extrinsic value (the degree to which predicted outcomes conform to my preferences) in simulating goal-directed exploration of a novel maze.

In summary, active inference casts perception as optimising beliefs about causes of sensory samples that minimise surprise (i.e., free energy) and action in terms of policies that minimise uncertainty (i.e., expected free energy). Expected free energy contains the right mixture of novelty, salience and prior preferences that constitute (Bayes) optimal beliefs about policies, which specify action (see Table 2 for a summary of the implicit resolution of surprise and uncertainty). Clearly, to evaluate posterior beliefs it is necessary to have a model of how states and parameters conspire to generate outcomes. So what does these models look like?

**Table 2:**
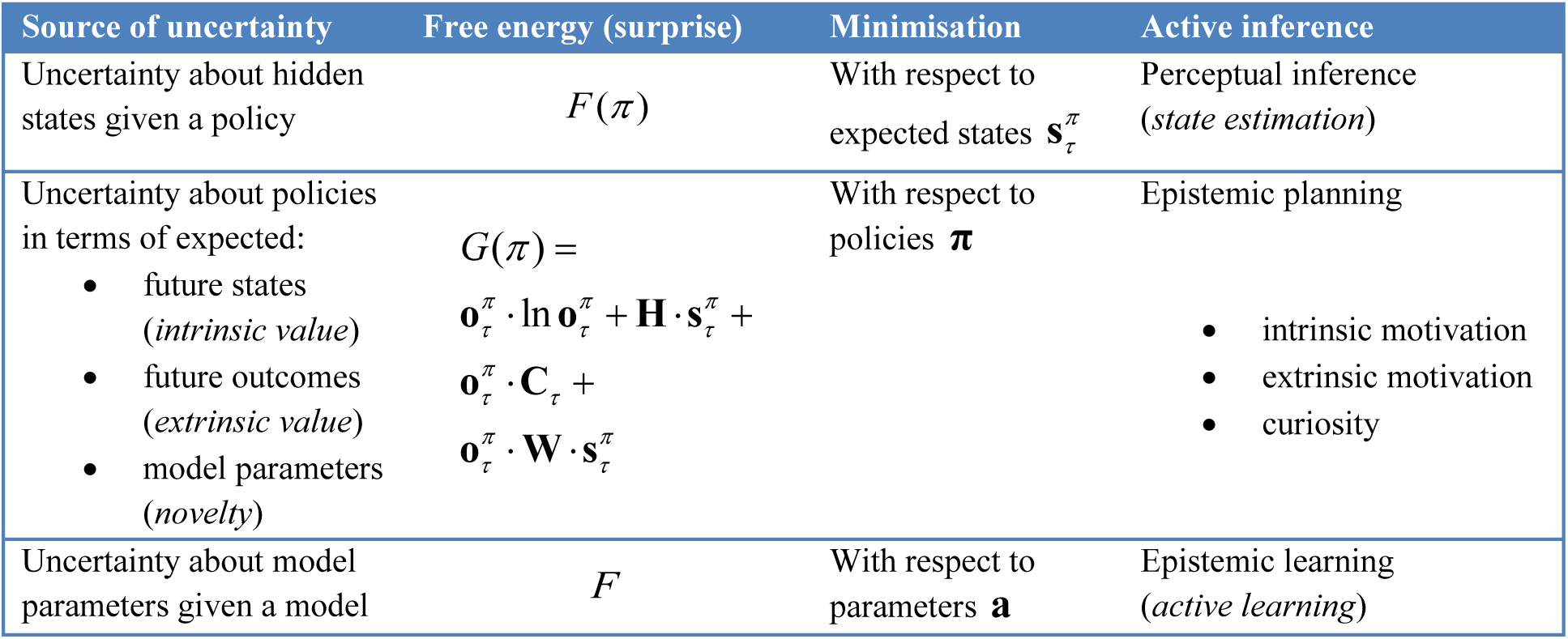
sources of uncertainty and the behaviours entailed by its minimisation; i.e., resolution of uncertainty through approximate Bayesian inference.

| Source of uncertainty | Free energy (surprise) | Minimisation | Active inference |
| --- | --- | --- | --- |
| Uncertainty about hidden states given a policy | $F(\pi)$ | With respect to expected states $\mathbf{s}_\tau^\pi$ | Perceptual inference (*state estimation*) |
| Uncertainty about policies in terms of expected:<br>future states (*intrinsic value*)<br>future outcomes (*extrinsic value*)<br>model parameters (*novelty*) | $G(\pi) =$<br>$\mathbf{o}_\tau^\pi \cdot \ln \mathbf{o}_\tau^\pi + \mathbf{H} \cdot \mathbf{s}_\tau^\pi +$<br>$\mathbf{o}_\tau^\pi \cdot \mathbf{C}_\tau +$<br>$\mathbf{o}_\tau^\pi \cdot \mathbf{W} \cdot \mathbf{s}_\tau^\pi$ | With respect to policies $\pi$ | Epistemic planning<br>intrinsic motivation<br>extrinsic motivation<br>curiosity |
| Uncertainty about model parameters given a model | $F$ | With respect to parameters $\mathbf{a}$ | Epistemic learning (*active learning*) |

### Generative models and Markov decision processes

Figure 1 provides a definition of a generic model that can be applied to most (discrete state space) scenarios. The particular form of the generative model used in this paper will be described in greater detail in the next section. In brief, a generative model is necessary to optimise beliefs about hidden states of the world and subsequent behaviour. This model is a probabilistic specification of how sampled outcomes are generated. For the sorts of Markov decision problems usually considered, it is sufficient to distinguish among four sorts of hidden or latent causes. These are hidden *states* generating outcomes, where transitions among hidden states are specified by a *policy*. This means there are two sorts of hidden states; namely, states of the world and the policies currently being pursued. As described above, beliefs over policies are proportional to the expected free energy or uncertainty under each policy. The constant of proportionality constitutes the third unknown; namely, the *precision* of beliefs about policies. This plays an interesting role in encoding the confidence in beliefs about the policies in play. It plays the same role as a softmax or inverse temperature parameter in classical softmax response rules and related formulations (Daw et al., 2011). Finally, the fourth unknown quantities are the *parameters* of the model. These correspond to matrices that specify the likelihood and (empirical) priors of the model. The first (*likelihood:* A) matrices encode the probability of outcomes under each hidden state, while the probability transition (*empirical prior:* B) matrices encode the probability of a subsequent state, given the current state. Crucially, there is a separate transition matrix for each allowable action, where a sequence of transitions is determined by the sequence of actions or policy. Prior beliefs about allowable policies depend on (precision weighted) expected free energy, which depend upon prior preferences over outcomes (*prior cost:* C), for each outcome modality over time. Finally, there are priors over the initial state (*initial priors*: D). When the parameters are unknown, they are usually modelled as Dirichlet distributions over the corresponding (likelihood and transition) probabilities. In other words, the underlying *concentration parameters* are essentially the frequency or number of times a particular outcome or state is generated from the hidden state in question. In what follows, we will only consider uncertainty about the likelihood. In other words, we will assume the agent knows the immediate consequences of action.

**Figure 1.**
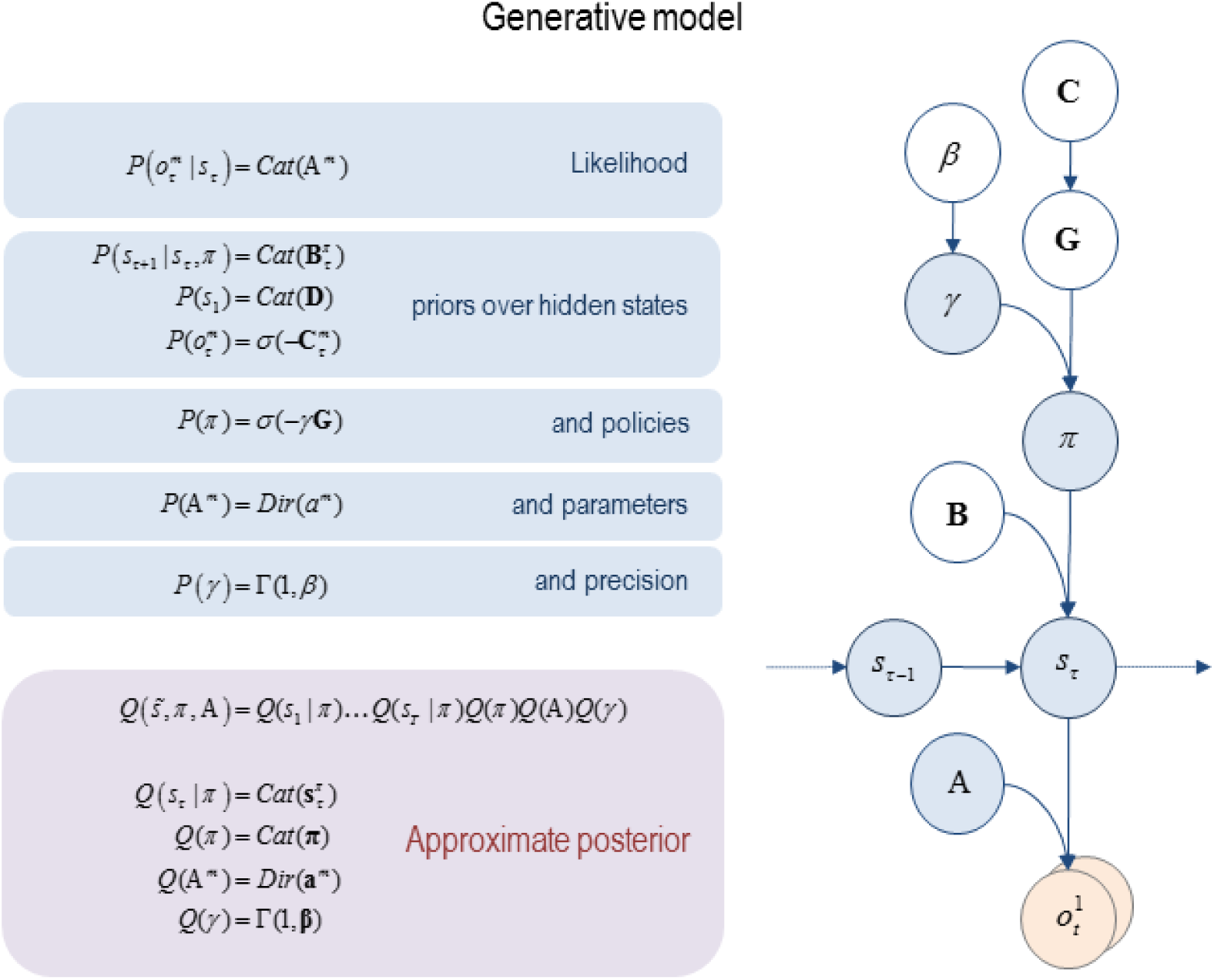
generative model and (approximate) posterior. A generative model specifies the joint probability of outcomes or consequences and their (latent or hidden) causes. Usually, the model is expressed in terms of a *likelihood* (the probability of consequences given causes) and priors over causes. When a prior depends upon a random variable it is called an *empirical prior*. Here, the likelihood is specified by matrices A whose components are the probability of an outcome under each hidden state. The empirical priors in this instance pertain to transitions among hidden states **B** that depend upon action, where actions are determined probabilistically in terms of policies (sequences of actions denoted by π). The key aspect of this generative model is that policies are more probable *a priori* if they minimise the (path integral of) expected free energy **G.** Bayesian model inversion refers to the inverse mapping from consequences to causes; i.e., estimating the hidden states and other variables that cause outcomes. In variational Bayesian inversion, one has to specify the form of an approximate posterior distribution, which is provided in the lower panel. This particular form uses a mean field approximation, in which posterior beliefs are approximated by the product of marginal distributions over unknown quantities. Here, a mean field approximation is applied both posterior beliefs at different points in time, policies, parameters and precision. See the main text and Table 2 for a detailed explanation of the variables. The insert shows a graphical representation of the dependencies implied by the equations on the right.

### Belief propagation in the brain

Equipped with this generative model, we can now derive update equations that minimise free energy. If this is done with careful respect for neurobiological constraints on the implicit Bayesian belief updating, one can derive a relatively straightforward process theory (Friston et al., 2015, Friston et al., 2016). The ensuing Bayesian belief updates are summarised in Figure 2. A detailed description of the belief update equations – and how they might be implemented in the brain – can be found in (Friston et al., 2017a). A complementary treatment that focuses on learning model parameters can be found in (Friston et al., 2016). The contribution of this paper is the form of the prior beliefs that underwrite policy selection. We therefore focus on these priors and how they inform policy selection through expected free energy. Appendix 2 derives the form of the subsequent belief updates (shown in Figure 2) for interested readers.

**Figure 2.**
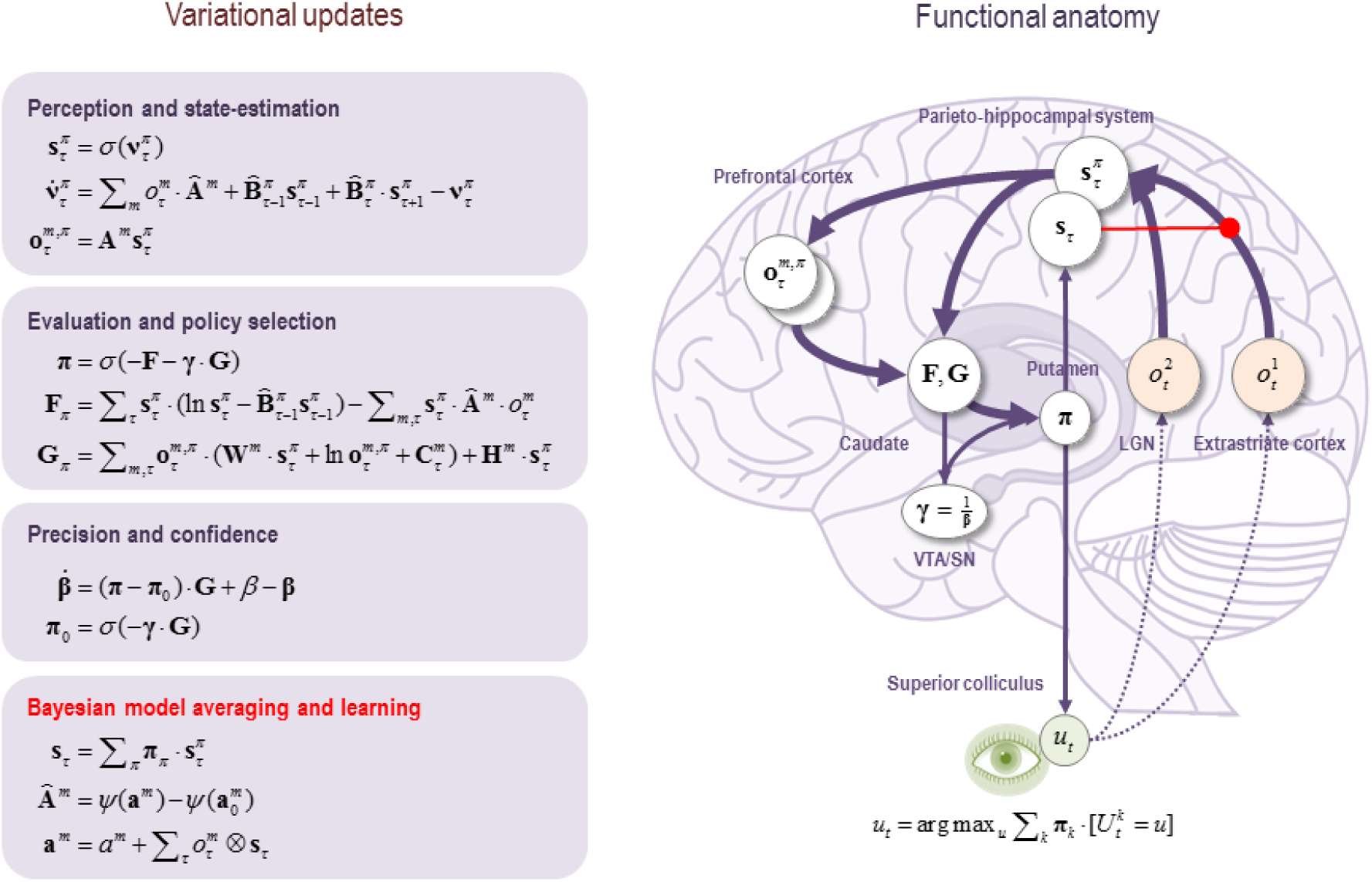
Schematic overview of belief updating. the left panel lists the belief updates mediating perception, policy selection, precision and learning; while the left panel assigns the updates to various brain areas. This attribution is purely schematic and serves to illustrate an implicit functional anatomy. Here, we have assigned observed outcomes to representations in the Pontine-geniculate occipital-system; with visual (*what)* modalities entering an extrastriate stream and proprioceptive (*where)* modalities originating from the lateral geniculate nucleus (LGN) via the superficial layers of the superior colliculus. Hidden states encoding location have been associated with the hippocampal formation and association (parietal) cortex. The evaluation of policies, in terms of their (expected) free energy, has been placed in the caudate. Expectations about policies – assigned to the putamen – are used to create Bayesian model averages of future outcomes (e.g., in the frontal or parietal cortex). In addition, expected policies specify the most likely action (e.g., via the deep layers of the superior colliculus). Finally, the precision of beliefs about – confidence in – policies rests on updates to expected precision that have been assigned to the central tegmental area or substantia nigra (VTA/SN). The arrows denote message passing among the sufficient statistics of each marginal as might be mediated by extrinsic connections in the brain. The red arrow indicates activity dependent plasticity. *Cat* and *Dir* referred to categorical and Dirichlet distributions respectively. Please see the appendix and Table 2 for an explanation of the equations and variables.

In brief, a belief propagation scheme is used to update the expected hidden states using a gradient descent on free energy. Crucially, posterior beliefs are over states from the beginning to the end of a trial or sequence of actions. This means that belief updating involves expectations about the past and future; enabling both prediction and postdiction separately under each policy. The second equation in Figure 2 (*perception and state estimation*) is an ordinary differential equation describing how expected hidden state is updated. The form of this equation – that falls out naturally from the form of the generative model – has a nice biological interpretation: this is because the updating involves the rate of change of a log expectation that is a sigmoid (softmax) function of log expectations (plus a decay term). This means that we can associate log expectations with depolarisation and implicit message passing with neuronal firing rates (that are a sigmoid activation function of depolarisation). In turn, this allows one to simulate electrophysiological responses in terms of fluctuations in log expectations.

These fluctuations occur at a number of timescales. At the fastest timescale they correspond to optimisation as the updates converge on a free energy minimum. This occurs following every new observation that, we assume, is sampled every 256 ms or so. These expectations are reset after a sequence of observations that we will refer to as a *trial*. In other words, a trial comprises a sequence of *epochs* in which an action is taken and a new outcome is observed. The length of a trial corresponds to the depth or horizon of the policies entertained by the agent. In what follows, we will use policies of two actions (that correspond to eye movements) and will call a two-action sequence a *trial* or *sub-path*.

Expectations about policies rest upon the posterior precision and expected free energy. Expected free energy in turn depends upon the expected states under the policy in question. This is a softmax function of expected free energy with a precision or inverse temperature parameter producing a conventional softmax response rule. Note that this form of probabilistic response emerges naturally from minimising variational free energy. The precision updates are effectively driven by the difference between the expected free energy over policies, relative to the equivalent expected free energy prior to observing outcomes. This is closely related to reward prediction error formulations and speaks to the similarity between precision updates and dopaminergic responses that we will appeal to later (Schultz et al., 2008, Friston et al., 2014, FitzGerald et al., 2015). Finally, the updates for the likelihood (concentration) parameters correspond to associative plasticity with trace and decay terms as discussed elsewhere (FitzGerald et al., 2015, Friston et al., 2016).

In short, we have a set of simple update rules for the four unknown quantities (namely, hidden states, policies, precision and parameters) that provide a process theory for state estimation, policy selection, confidence and learning respectively. Note that these equations are completely generic. In other words, they are exactly the same equations used in all previous illustrations of active inference under Markov decision process models: e.g., (Friston et al., 2014, FitzGerald et al., 2015, Friston et al., 2015, Friston et al., 2016, Mirza et al., 2016). In the next section, we will see examples of this updating cast in terms of simulated neuronal and behavioural responses. However, before simulating these responses, it is necessary to specify the precise form of the generative model.

## A generative model for planning

This section describes the particular form of the generative model – in terms of its parameters, hidden states and policies – that will be used in the remainder of this paper. This model captures the bare essentials of a maze foraging task under novelty or uncertainty. In brief, the (synthetic) subject sees a maze specified on an 8 x 8 grid. The subject initially fixates on a starting location and then has to navigate to a target location deep within the maze. Notice that we are simulating a maze that can be interrogated with a visual search – as opposed to simulating a physical maze of the sort that a rat would explore. This is because we hope to use this model to explain empirical responses from (human) subjects performing the task. Having said this, we imposed constraints on the sampling of the maze so that it was isomorphic with a (myopic) rat exploring a physical maze. This was implemented by restricting movements or actions to single steps in four directions (or remaining at the same location). Furthermore, the sensory (visual) outcomes were limited: the subject could only see whether the current location was accessible (*open* – white) or not or (*closed* – black). These constraints meant that – in the absence of any knowledge about the maze that has been accrued through previous learning or experience – the subject had to forage for local information to build an internal model of the maze structure, using short sequences of saccadic eye movements. Crucially, the agent could only entertain shallow policies of two moves. In other words, they could only consider (25) policies corresponding to all combinations of five actions (*up*, *down*, *left*, *right*, or *stay*). This is an important constraint that precluded an exhaustive (deep) search to identify the best policy for reaching the goal. In effect, this enforces a chunking of the problem into subgoals entailed by prior beliefs about outcomes or preferences.

In addition to visual input, we also equipped agents with positional information; namely, the current location that they occupied. This meant that there were two outcome modalities: *what* (open versus closed) and *where* (among 64 locations). The generative model of these outcomes was very simple: the hidden states corresponded to the location (with 64 possibilities). The likelihood mapping from location to outcomes comprised two A matrices, one for each outcome modality. The first simply specified the probability of observing open versus closed at each location, while the second was an identity mapping returning the veridical location for each hidden state. The (empirical) prior transition probabilities were encoded in five B the matrices. Again, these were very simple and moved the hidden (*where*) states to the appropriate neighbouring location, unless the action was *stay* or transgressed the boundary of the maze – in which case the location did not change.

This generative model can be used in two modes. We can either assume that the subject has had a lot of experience with a particular maze and has accrued sufficient evidence to learn the mapping between location and outcome. In other words, knowledge about the maze is encoded in terms of what would be seen at each location. The subject can use this information to plan or navigate through the maze from the starting location to a target as described below – providing we instil the necessary prior beliefs. Alternatively, we could assume that the maze is novel. In this context, the (concentration) parameters of the likelihood mapping to *what* outcomes will be uniformly very small for all locations (we used 1/8). In this instance, the subject has to first learn the maze before she can perform the task.

Figure 3, shows the learning of the maze over 64 eye movements, using high values (128) of the concentration parameters for the mapping between location and the *where* modality. Learning is shown in terms of the accumulated concentration parameters for the *what* modality that are garnered by epistemic foraging. The key aspect of this behaviour is that the movements are driven by novelty, successively exploring unexplored regimes of the maze until all epistemic value or information gain has been consumed. In other words, once a location has been visited it is no longer novel or attractive, thereby rendering the probability that it will be sampled again less likely: c.f., inhibition of return (Wang and Klein, 2010). This epistemic foraging is driven entirely by the novelty afforded by ignorance about the first (*what*) modality. In other words, noise or stochastic fluctuations are unnecessary for generating explorative behaviour. The reason that exploration was driven entirely by novelty is that there is no uncertainty about hidden states (given the precise and ambiguous outcomes in the *where* modality – and the fact that there were no prior preferences to constitute place preferences).

**Figure 3.**
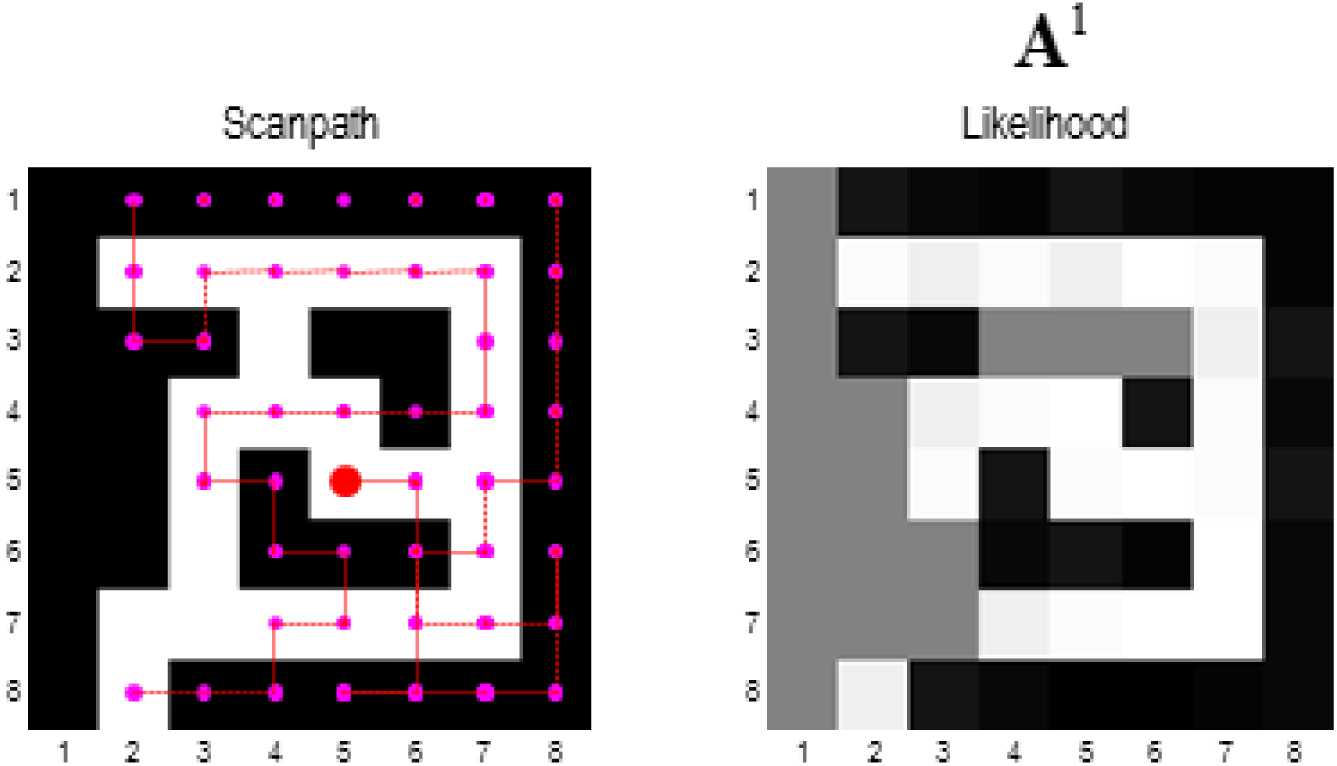
explorative, epistemic behaviour. Left panel: This figure reports the results of epistemic exploration for 32 (two move) trials (e.g., 64 saccadic eye movements). The maze shown in terms of closed (black) and open (white) locations. The magenta dots and lines correspond to the chosen path, while the large red dot denotes the final location. The agent starts (in this maze) at the entrance on the lower left. The key thing to observe in these results is that the trajectory very seldom repeats or crosses itself. This affords a very efficient search of state space, resolving ignorance about the consequences of occupying a particular location (in terms of the first – *what* – outcome; black vs. White). **Right panel**: this figure reports the likelihood of observing an open state (white), from each location, according to the concentration parameters of the likelihood matrix that have been accumulated during expiration (for the first – *what* – outcome modality). At the end of search, the posterior expectations change from 50% (grey) to high or low (white or black) in, and only in, those locations that have been visited. The underlying concentration parameters effectively remember what has been learned or accumulated during exploration – and can be used for planning, given a particular task set (as illustrated in the Figure 5).

Figure 4 shows the simulated electrophysiological responses during the exploration above. These predicted responses are based upon the optimisation of expected hidden states and precision, described by the differential equations that mediate belief propagation (see Figure 2). It is these sorts of responses that are underwritten by the process theory on offer. The results in Figure 4 serve to illustrate the potential for generating predictions of behavioural, electrophysiological and dopaminergic responses that could be used in empirical studies. See (Schwartenbeck et al., 2015) for an example of using these sorts of simulated responses in computational fMRI.

**Figure 4.**
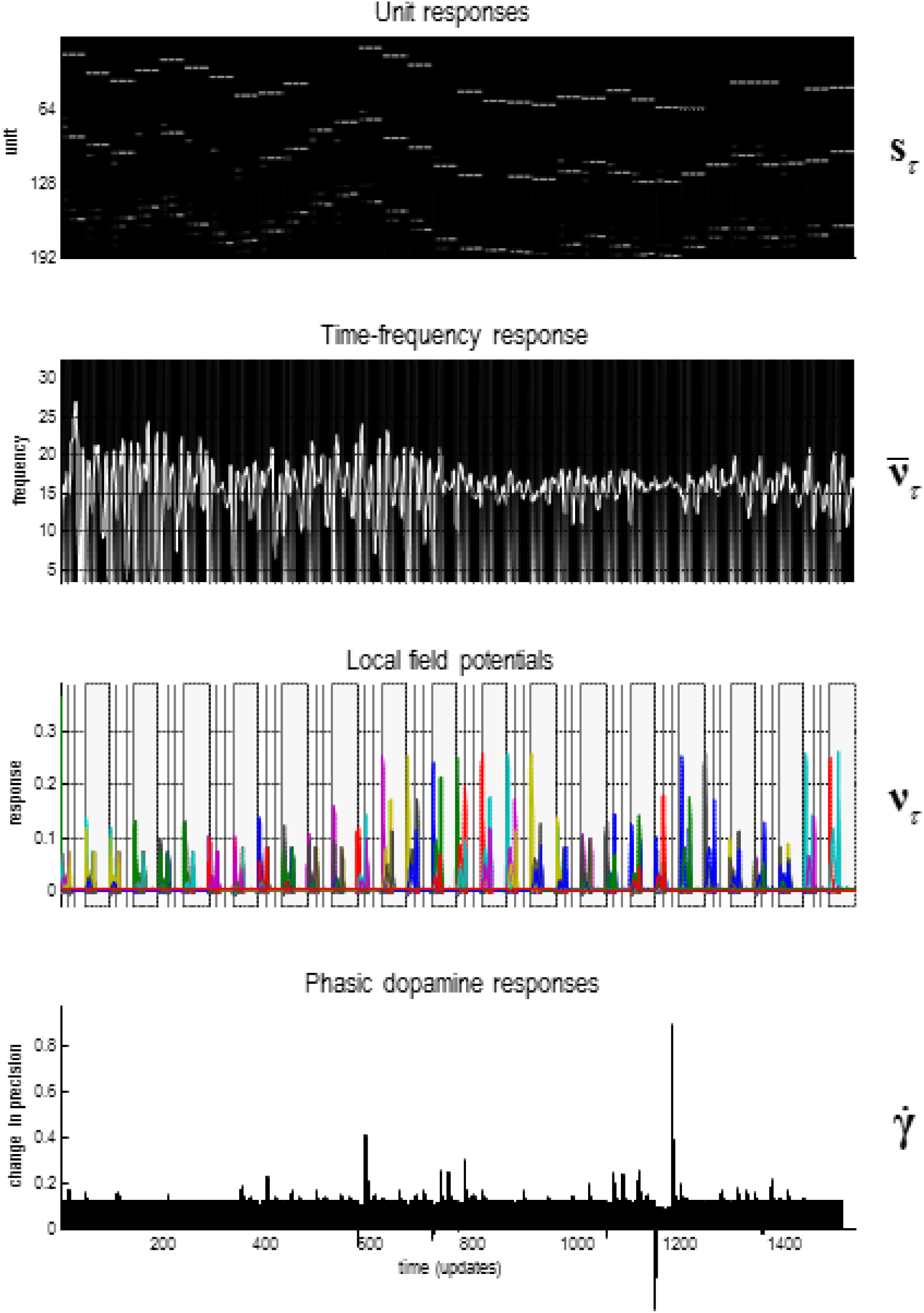
Simulated electrophysiological responses during exploration: this figure reports the simulated electrophysiological responses during the epistemic search of the previous figure. **Upper panel**: this panel shows the activity (firing rate) of units encoding the expected location – over 32 trials – in image (raster) format. There are 192 = 64 x 3 units for each of the 64 locations over the three epochs between two saccades that constitute a trial. These responses are organised such that the upper rows encode the probability of alternative states in the first epoch, with subsequent epochs in lower rows. The simulated local field potentials for these units (i.e., log state prediction error) are shown in the middle panels. **Second panel**: this panel shows the response of the first hidden state unit (white line) after filtering at 4 Hz, superimposed upon a time-frequency decomposition of the local field potential (averaged over all units). The key observation here is that depolarisation in the 4 Hz range coincides with induced responses; including gamma activity. **Third panel**: these are the simulated local field potentials (i.e. depolarisation) for all (192) hidden state units (coloured lines). Note how visiting different locations evokes responses in distinct units of varying magnitude. Alternating trials (of two movements) are highlighted with grey bars. **Lower panel**: this panel illustrates simulated dopamine responses in terms of a mixture of precision and its rate of change (see Figure 2). There phasic fluctuations reflect changes in precision or confidence based upon the mismatch between the free energy before and after observing outcomes (see Figure 2).

The upper panel shows the simulated firing rates of units encoding the expected location over the three epochs that surround the two eye movements that constitute successive trials. The fluctuations in transmembrane potential that drive these firing rates can be used to simulate induced responses (second panel) or evoked responses (third panel). The induced or time frequency responses over all units (second panel) are interesting from the point of view of theta-gamma coupling in the hippocampus during exploration (Dragoi and Buzsaki, 2006, Colgin et al., 2009; Lisman and Redish, 2009, Jezek et al., 2011, Buzsaki and Moser, 2013). This coupling arises naturally as the fast (gamma) optimisation of posterior expectations is entrained by a slow (theta) sampling of the environment (Friston and Buzsaki, 2016). Finally, the lower panel shows simulated dopaminergic responses in terms of the rate of change of precision (plus an offset). Note how posterior precision fluctuates more markedly as the maze becomes more familiar and the subject becomes more confident about what she is doing. We will take a closer look at predicted electrophysiological responses in the last section. First, we consider how goal seeking emerges when we add prior preferences.

### Prior preferences, constraints and goals

To simulate navigation *per se*, we need to consider the prior beliefs about outcomes that engender goal-directed behaviour. In this paper, we will adopt a particular scheme; noting that many other plausible priors (i.e., heuristics) could have been used. We embrace this plurality because, ultimately, we want to adjudicate amongst different priors when trying to explain the empirical responses of real subjects. However, here, we will focus on one straightforward but efficient formulation.

The prior preferences that lead to purposeful navigation – i.e. task set – can be specified purely in terms of prior preferences over location outcome. These reflect the subject’s beliefs about what are plausible and implausible outcomes and the constraints under which she operates. To accommodate these constraints, we used the following heuristic: namely, that a subject believes that she will occupy locations that are the most accessible from the target. If these prior preferences are updated after each trial, the subject will inevitably end up at the target location. To evaluate the accessibility of each location from the target location, the agent can simply evaluate the probability it will be occupied under allowable state transitions from the target location. Formally, this can be described as follows:

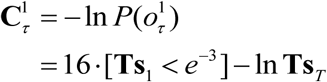

The first term assigns a high cost to any location that is occupied with a small probability, when starting from the initial location. The second term corresponds to the (negative) log probability a given state is occupied, when starting from the target location (encoded by **s**_*T*_). Prior beliefs about allowable transitions **T** are based upon posterior beliefs about the structure of the maze; namely, whether any location is open or closed.

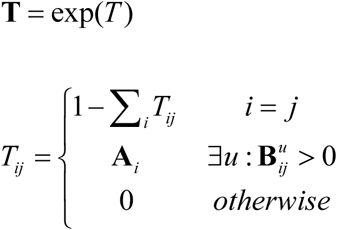

The probability transition matrix **T** plays the role of a Green’s function based upon the graph Laplacian *T*. Intuitively, **T** encodes the probability that any state will be occupied following diffusion from any other state *during one time step* – and the graph Laplacian encodes allowable paths based upon the posterior beliefs about the maze. Specifically, the graph Laplacian comprises the posterior probability that a state is *open* and can be reached by an action from another state.

This particular heuristic was chosen to formalise the intuition that we decompose distal goals into intermediate subgoals and, in particular, attainable subgoals under an admixture of constraints. In other words, we tend to select those states that can be reached that can also be reached from the target state: see discussion and (Dijkstra, 1959). This suggests we contextualise our subgoals using knowledge about the ultimate goal so that the (forward and backward) passes through some problem space ‘meet in the middle’. Note the formal similarity between backwards induction and optimisation of state action policies under the Bellman optimality principle (Bellman, 1952) and the diffusion heuristic above. However, this heuristic goes a bit further and augments the implicit (extrinsic) value function of location with a reachability cost. This acknowledges the fact that, in active inference, agents are already prospective in their policy selection; even if the time horizon of these policies is not sufficient to reach the ultimate goal. Technically, active inference for Markov decision processes entails sequential policy optimisation – as opposed to optimising state-action policies. Having said this, state-action policies can be learned as habits under active inference, provided they are fit for purpose (Friston et al., 2016). Here, we effectively use a simple form of backwards induction to contextualise sequential policies with a limited horizon.

As noted above, there are probably many schemes or heuristics that one could use. The diffusion heuristic appears to be remarkably efficient and is sufficient for our purposes; namely, to show how prior preferences about (*where*) outcomes can subsume all the prior beliefs necessary for instantiating a task set necessary for navigation. Figure 5 shows the results of a typical navigation when the maze is familiar; i.e., it is known *a priori*. The middle panels show the prior preferences (extrinsic value) over locations during navigation. Note how the locations with greatest extrinsic value effectively lead the agent towards the target location; thereby playing the role of subgoals. This simulation used concentration parameters – encoding the structure of the maze – of 128. This corresponds to 128 exposures to each location. One might now ask how navigation depends upon experience.

**Figure 5.**
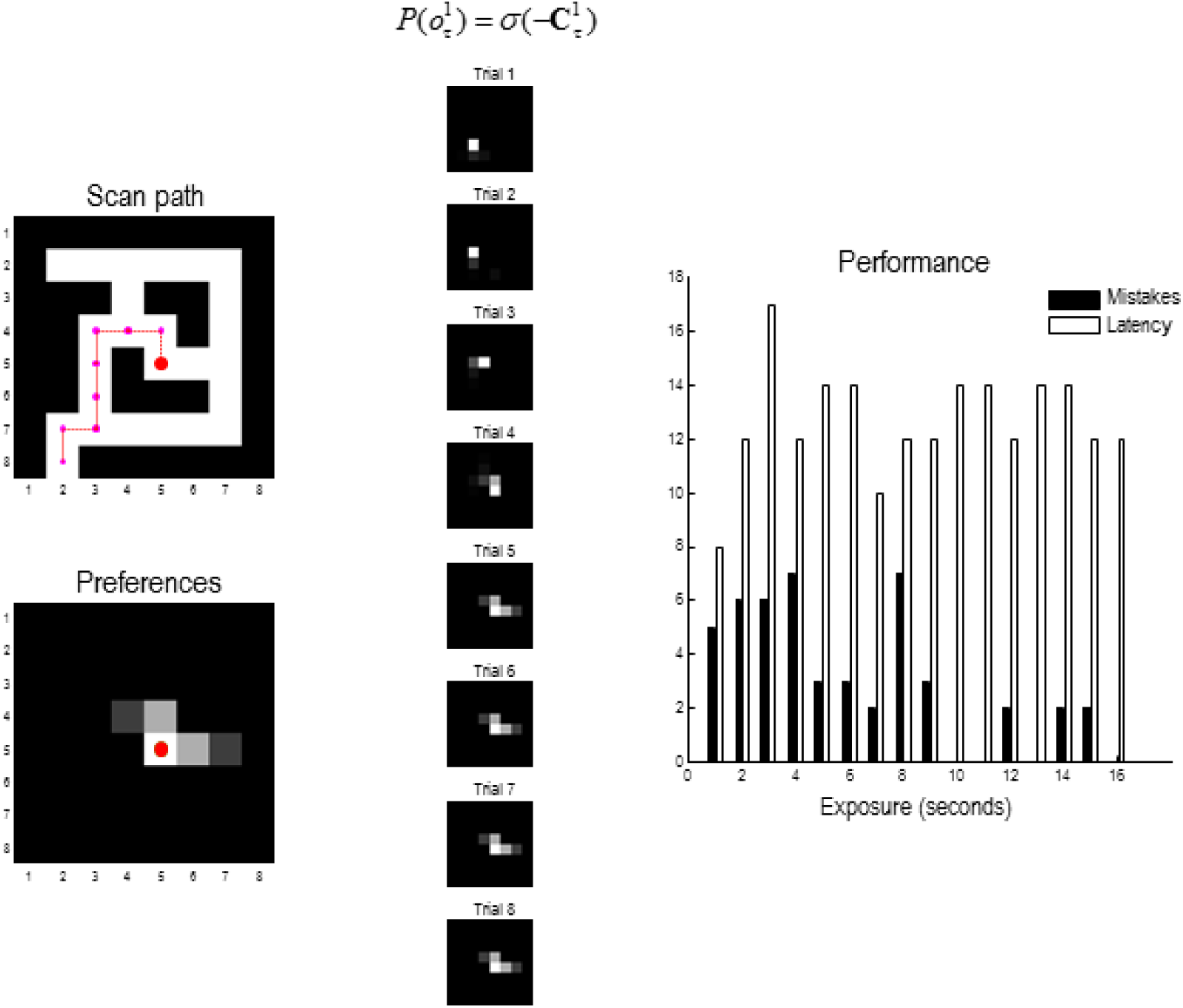
planning and navigation. this figure shows the results of navigating to a target under a task set (i.e., prior preferences), after the maze has been learned (with the concentration parameters of 128). These prior preferences render the closed (black) locations surprising and they are therefore avoided. Furthermore, the agent believes that it will move to locations that are successively closer to the target – as encoded by subgoals. **Left panels**: the upper panel shows the chosen trajectory that takes the shortest path to the target, using the same format as Figure 3. The lower panel shows the final location and prior preferences in terms of prior probabilities. At this point, the start and end locations are identical – and the most attractive location is the target itself. As earlier points in navigation, the most attractive point is within the horizon of allowable policies. **Middle panels**: these show the prior preferences over eight successive trials (16 eye movements), using the same format as above. The preferred locations play the role of context sensitive subgoals; in the sense that subgoals lie within the horizon of the (short-term) policies entertained by the agent – and effectively act as a ‘carrot’ leading the agent to the target location. **Right panel**: these report the planning or goal directed performance based upon partially observed mazes, using the simulations reported in Figure 3. In other words, we assessed performance in terms of the number of moves before the target is acquired (*latency*) and the number of closed regions or disallowed locations visited *en route (mistakes*). These performance metrics were assessed during the accumulation of concentration parameters. This corresponds to the sort of performance one would expect to see if a subject was exposed to the maze for increasing durations (here, from one to 16 seconds of simulated time), before being asked to return to the start location and navigate to a target that is subsequently revealed.

The right panel of Figure 5 shows navigation performance (in terms of the time taken to secure the goal and transgressions into *closed* locations) as a function of familiarity with the maze. Familiarity was simulated using the accumulated concentration parameters from the exploratory simulations above. In other words, we effectively expose the subject to the maze for increasing durations of time (2 to 16 seconds of simulated – and roughly computer – time). The degree to which familiarity supported task performance was then assessed by recording the path taken to the target when starting from the initial location. These results show that the subject was able to navigate to the goal, without making any mistakes, after about 16 seconds of exploration (no mistakes were made after 16 seconds in this example). Note that the time taken to reach the target is paradoxically the shortest when the maze is unfamiliar (i.e. on the first exposure), because the subject took shortcuts along illegal paths. In the final section, we ask what would happen when behaviour is driven by both epistemic and pragmatic value at the same time.

## Goal directed exploration

Finally, we turn to the integration of intrinsic (epistemic) and extrinsic (pragmatic) value by equipping the agent with goal-directed prior beliefs (i.e., task set) during the epistemic foraging. The main purpose of this simulation is to show active inference dissolves the exploration–exploitation dilemma by absorbing extrinsic and intrinsic imperatives into a single objective function (i.e., expected free energy). Heuristically, the ensuing behaviour is initially driven by epistemic imperatives until sufficient uncertainty has been resolved to realise the pragmatic or extrinsic imperatives. This is precisely what we see in the current setup.

Figure 6 shows the results of foraging for information under the goal directed prior beliefs above. Here, we see that the exploration is now (goal) directed. In other words, as soon as there is sufficient information about the structure of the maze, it is used to constrain epistemic foraging, until the target is reached. In these simulations, the subject navigated to the target location during four successive searches that were limited to eight trials or 16 eye movements. It can be seen that perfect (shortest path) performance is attained by the fourth attempt. Prior to this, there are excursions into closed locations. Interestingly, when the maze is still novel, curiosity gets the better of the subject and the target location is largely ignored in favour of resolving uncertainty about nearby locations.

**Figure 6.**
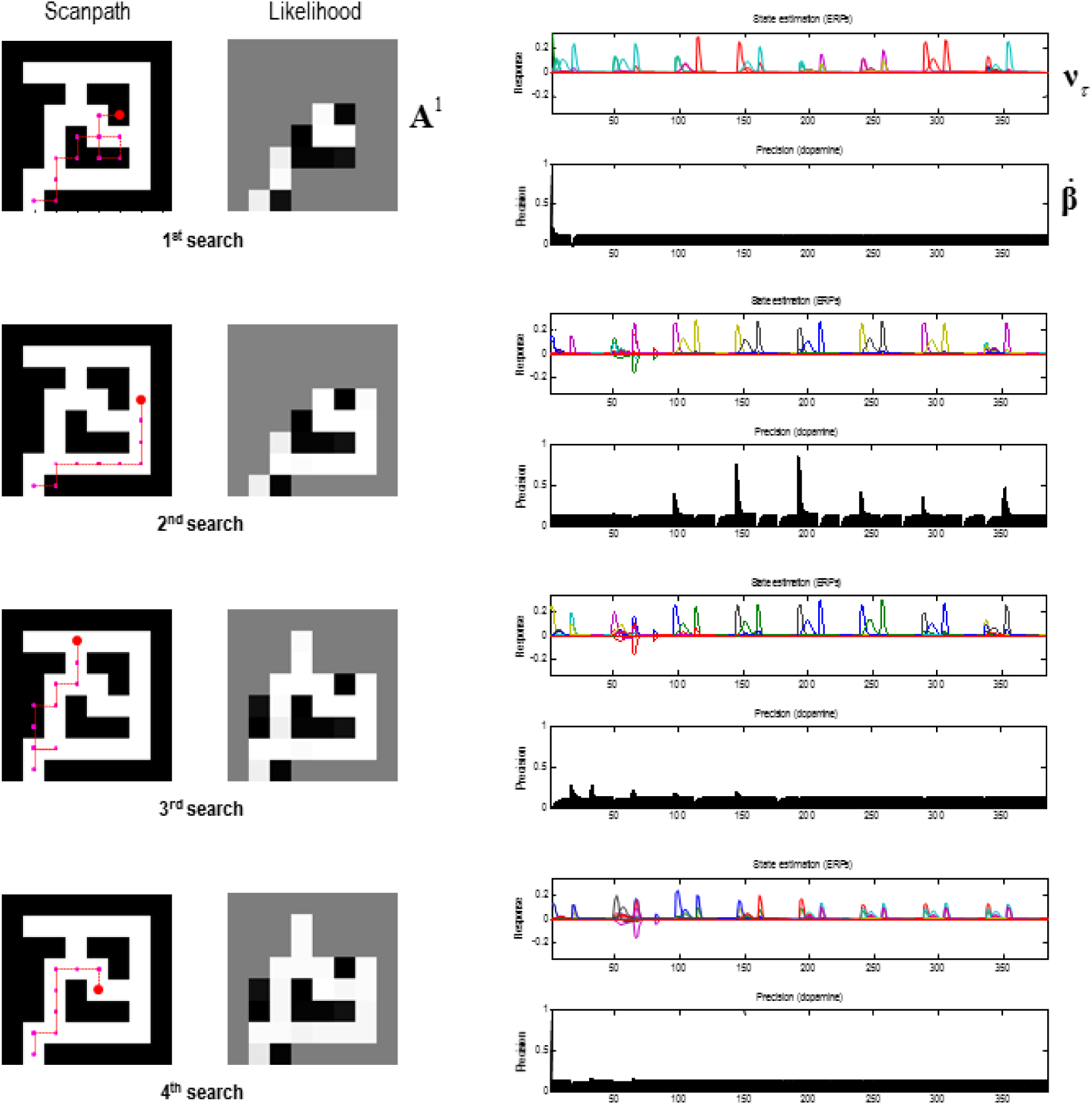
goal-directed exploration: this figure illustrates behavioural, mnemonic and electrophysiological responses over four searches, each comprising 8 trials (or 16 eye movements). Crucially, the agent started with a novel maze but was equipped with a task set in terms of prior preferences leading to the goal directed navigation of the previous figure. Each row of panels corresponds to a successive search. **Left panels**: these report the path chosen (left) and posterior expectations of the likelihood mapping (right) as evidence is accumulated. However, here, the epistemic search is constrained by prior preferences that attract the target. This attraction is not complete and there are examples where epistemic value (i.e., novelty of a nearby location) overwhelms the pragmatic value of the target location – and the subject gives way to curiosity. However, having said that, the subject never wanders far from the shortest path to the target, which she acquires optimally after the fourth attempt. **Right panels**: these show the corresponding evoked responses or simulated depolarisation in state units (upper panels) and the corresponding changes in expected precision that simulate dopaminergic responses (lower panels). The interesting observation here is the progressive attenuation of evoked responses in the state units as the subject becomes more familiar with the maze. Interestingly, simulated dopaminergic responses suggest that the largest phasic increases in confidence (i.e., a greater than expected value) are seen at intermediate points of familiarity; while the subject is learning the constraints on her goal directed behaviour. For example, there are only phasic decreases in the first search, while phasic increases are limited to subsequent searches.

The right hand panels of Figure 6 show the corresponding simulated physiological responses, using the format of Figure 4. In this example, we see systematic and progressive changes in (simulated) electrophysiological and dopaminergic responses. The latter are particularly interesting and reflect the fact that as the subject engages with novel opportunities, she resolves uncertainty and thereby suppresses fluctuations in precision or confidence. To our knowledge, this has not been addressed empirically and represents a prediction of the current simulations. Namely, one would expect to see more phasic dopamine responses (or fMRI responses in the target regions of the dopaminergic projections) during the initial exploration of a maze, relative to later periods that may be more exploitative in nature. This is a somewhat paradoxical prediction that, in principle, could be confirmed with human subjects and fMRI: c.f., (Bunzeck and Duzel, 2006, D’Ardenne et al., 2008, Schwartenbeck et al., 2015).

### Place cells or path cells or both?

As noted in the introduction, the purpose of this work was to formulate spatial navigation in terms of active inference, using the same scheme that has been used to model several other decision-making, cognitive and perceptual paradigms. This scheme has a fairly well established process theory that allows one to make specific predictions about electrophysiological and psychophysical responses. There are many avenues that one could pursue in light of the simulations described in this paper. For example, the encoding of hidden states in terms of location leads naturally to a formulation in terms of place cells and the attendant spatiotemporal encoding of trajectories through space. In this setting, one could associate the encoding of policies with direction selective neuronal responses (Taube, 2007) – and consider how simulated place cell activity (expectations about hidden states) depend upon direction cells (i.e., expectations about policies) (Lisman and Redish, 2009). These predictions would speak against a simple (orthogonal) encoding of place and direction and would predict particular forms of joint peristimulus time histogram responses – that reflect the message passing between representations of state (i.e., place) and policy (i.e., directed trajectory). We will pursue this elsewhere and focus here on an even simpler insight afforded by the above simulations.

In virtue of inferring the best policy, in terms of its consequences for latent states of the world, there is a necessary encoding of future states under allowable policies. This brings something quite interesting to the table; namely, the encoding trajectories or paths through state space. In particular, it means that certain units will encode the location at the start of any sequence of movements – and will continue doing so until a subgoal is attained. Conversely, units that encode future states will only respond when there is clear evidence that the subgoal has been reached. This dissociation – between the predicted firing patterns of units encoding hidden states at the beginning and ends of sub-paths – means that there must be a continuum of place specificity that could present itself as place-cell-like activity of greater or lesser precision. In other words, if one subscribes to the neuronal process theories above, then place cell responses can be regarded as the limiting case of a more general encoding of local paths (Knierim et al., 2014, Friston and Buzsaki, 2016).

This is illustrated in Figure 7, which plots the activity of several units as a function of location in the maze (see middle panels). Crucially, cells encoding the initial location at the beginning of each subpath maintain their firing during the course of the path to the subgoal. Conversely, cells that encode hidden states towards the end of each sub-path have a spatial specificity, because they are only engaged when the location is reached. It is tempting to think of neuronal encoding in terms of ‘path cells’ that encode where the (synthetic) agent has been recently; such that a proportion of these path cells – encoding hidden states at the end of a local trajectory – become ‘place cells’ proper. If this form of spatiotemporal encoding is in play in real brains, it suggests that there may be more information in neuronal responses available for reconstructing spatial paths or trajectories than based solely on canonical place cell activity (Guger et al., 2011). This speaks to a characterisation of neuronal responses both in terms of the position of a rat and its local trajectory in temporal frames of reference that maybe anchored to subgoals (Pfeiffer and Foster, 2013). Clearly, this would be a challenging but interesting possibility that might nuance our understanding of spatiotemporal encoding; in particular, scheduling in temporal frames of reference that transcend our normal conception of the past and future (Eichenbaum, 2014).

**Figure 7.**
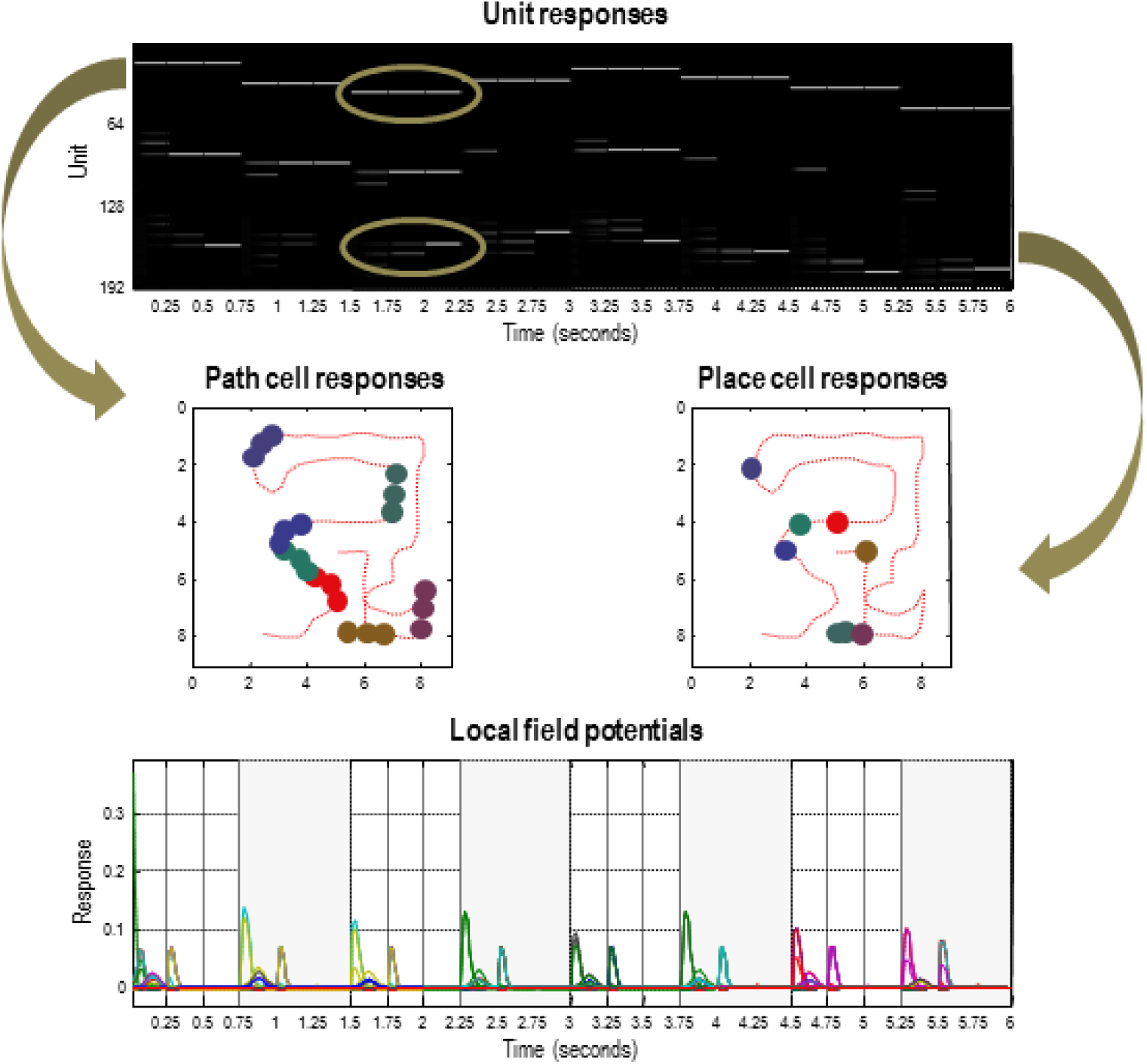
path and place cells: this figure revisits the simulation in Figure 4 but focusing on the first six seconds of exploration. As in Figure 4, the upper panel shows the simulated firing of the (192) units encoding expected hidden states, while the lower panel shows the accompanying local field potentials (obtained by band-pass filtering the neuronal activity in the upper panel). The key point made in this figure is that the first 64 units encode the location at the start of each local sequence of moves and maintain their firing until a subgoal has been reached. Conversely, the last 64 units encode the location at the end of the local sequence and therefore only fire *after* the accumulation of evidence that a subgoal has been reached. This leads to an asymmetry in the spatial temporal encoding of paths. In other words, first set of units fire during short trajectories or paths to each subgoal, while the last set only when a particular (subgoal) location has been reached. This asymmetry is highlighted by circles in the upper panel (for the third sub-path), which shows the first (upper) unit firing throughout the local sequence and the second (lower) unit firing only at the end. The resulting place preferences are illustrated in the middle panels; in terms of *path cell* (left panel) and *place cell* (right panel) responses. Here, we have indicated when the firing of selected units exceeds a threshold (of 0.8 Hz), as a function of location in the maze during exploration (the dotted red line). Each unit has been assigned a random colour. The key difference between path and place cell responses is immediately evident, path cells respond during short trajectories of paths through space, whereas place cell responses are elicited when, and only when, the corresponding place is visited.

## Discussion

In what follows, we consider related approaches and previous computational formulations of spatial planning. We then briefly consider the known functional neuroanatomy engaged by these sorts of tasks. This discussion is presented as a prelude to subsequent work that will use the current model to fit the behavioural and fMRI responses elicited by spatial planning in humans (Kaplan et al., 2017a).

### Relationship to Previous Work

Many of the assumptions entailed by the priors of the generative model in this paper are inherited from previous work using the principles of optimal control and dynamic (reinforcement) learning to understand navigation through (state) spaces and implicit deep tree searches. In terms of systems and cognitive neuroscience, reinforcement learning paradigms provide a nice taxonomy within which to place the current active inference formulation. Put simply, spatial navigation and, more generally planning, presents an intractable deep tree search into the future. There are at least six ways in which one can avoid full tree searches (Mehdi Keramati – personal communication). First, one can formulate the problem in terms of *habit learning*. That is, avoid planning altogether and rely upon state-action policies that are accrued in the usual way through caching the value of actions from any given state (Sutton and Barto, 1998). Some key theoretical papers in this setting include (Sutton and Barto, 1998, Daw et al., 2005, Keramati et al., 2011). Empirical evidence – for habits in the brain – derives from neuroimaging and the use of normative models based upon reinforcement learning (Daw et al., 2011, Lee and Keramati, 2017). Second, one can *prune* or limit the depth of the tree search. See (Huys et al., 2012) for a discussion of how a (Pavlovian) system could sculpt choices by pruning decision trees. A nuanced version of pruning involves planning until a certain depth and then switching to habitual value estimation at the leaves of the search tree; i.e., *plan until habit* (Keramati et al., 2016). This combines the efficiency of habit learning, yet still retains a degree of context sensitivity via planning. An alternative approach – *bidirectional planning* – rests upon parallel searches of decision trees; one search starting from the current (inferred) state and another from the goal state (Dijkstra, 1959). An alternative, known as *hierarchal decision-making*, involves planning on an abstract representation of a Markov decision process (known as the ‘*option’* framework in the hierarchal reinforcement literature). This is reviewed in (Botvinick et al., 2009), and enjoys a degree of empirical support (Ribas-Fernandes et al., 2011, Collins and Frank, 2016). Finally, *successor representation* involves caching successor states that can be reached from each state action pair (Dayan, 1993). These representations can be used to estimate the value of state action pairs when combined with the reward associated with each state. Again, there is some experimental evidence for this formulation (Momennejad et al., 2017, Russek et al., 2017).

These reinforcement learning approaches are formally distinct from active inference because they do not accommodate the effects of policies or action on belief states that may nuance optimal sequences of behaviour. This is a fundamental distinction that can be reduced to the following: active inference optimises a *functional of beliefs about states*; namely, the expected free energy above. This contrasts with reinforcement learning – and related approaches based upon the Bellman optimality principle – that try to optimise a *function of states per se* – as opposed to beliefs about states (Friston et al., 2015). Having said this, there are many aspects of the above reinforcement learning formulations that we have appealed to in this paper. For example, there are elements of hierarchal decision-making and successor representations implicit in the deep temporal (generative) model that underlies inferences about policies and subsequent policy selection. Furthermore, bidirectional planning is, in a loose sense, implicit in the bidirectional message passing between active inference representations of the past and future: see also (Gershman, 2017). As in pruning approaches, this bidirectional aspect is kept to a manageable size by the induction of subgoals that allow for a chunking or decomposition of the tree search. It remains an interesting and challenging exercise to migrate reinforcement learning schemes into the world of belief states (e.g., partially observed Markov decision processes). The current treatment is an attempt to leverage these ideas in the setting of active inference.

The imperative to resolve epistemic uncertainty, under active inference, fits comfortably with recent work using variational inference in Bayesian neural networks to maximize information gain during exploration (Houthooft et al., 2016). Artificial curiosity in simulated agents is an essential aspect of spatial planning (Vigorito and Barto, 2010). Curious, uncertainty resolving behaviour arises in our scheme via the selection of policies that not only reduce uncertainty about hidden states of the world (i.e., salience) but also reduce ignorance about hidden contingencies encoded by the parameters of the agent’s generative model; i.e., novelty (Friston et al., 2017b). The ensuing resolution of uncertainty through information gain is exactly as articulated in terms of planning to be surprised: see (Sun et al., 2011b). In the current formulation, the information gain in question is a constituent of expected free energy; namely, the epistemic value that underwrites exploratory behaviour.

Our formulation contributes to an emerging literature on multi-step planning in novel environments. Here, we focused on a spatial planning task that involves a myopic agent making saccadic eye movements, while learning the quickest path to a goal location. Other studies have investigated planning in novel environments (McNamee et al., 2016) or puzzle-like tasks (Solway et al., 2014, Maisto et al., 2015). Despite these differences, all of these studies entail hierarchical problems that are solved by chunking action sequences (Fonollosa et al., 2015). In the context of planning, this chunking is known as subgoaling (van Dijk and Polani, 2011, Van Dijk and Polani, 2013, Maisto et al., 2015, Donnarumma et al., 2016); where the agent locates subgoals/bottlenecks *en route* to achieving a goal. Due our use of a graph Laplacian – in forming prior preferences – the imperative to reach subgoals emerges without any explicit marking of subgoals in the environment (e.g. explicitly informing the agent where a choice point is located). In other words, the agent behaves ‘as if it was securing a succession of subgoals; however, these subgoals are essentially phenomenological.

One prominent advantage of locating subgoals within an environment, regardless of whether a subgoal is spatial or abstract, is that it affords modularization of a state space; in the service of focusing on relevant sub-tasks (Solway and Botvinick, 2012, Solway et al., 2014, Solway and Botvinick, 2015). State-space modularization is particularly important during online spatial planning in novel environments (McNamee et al., 2016). Recent work suggests that the type of online state-space modularization captured by McNamee and colleagues (McNamee et al., 2016) might rely on the hippocampal formation and anterior prefrontal cortex – brain areas thought to signal the current location within a state space, relative to a goal (Kaplan et al., 2017b).

In neurobiology, a well-studied modularization of space at the level of single neurons occurs in grid cells located in the dorsomedial entorhinal cortex (Hafting et al., 2005). Some modelling work has already made substantial progress in that direction – by showing that populations of grid cells can guide goal-directed navigation (Erdem and Hasselmo, 2012, Bush et al., 2015). Systems level models have been proposed to address complex spatial planning behaviour; for example, Martinet and colleagues (Martinet et al., 2011) modelled a prefrontal-hippocampal network that could perform multi-level spatial processing, encode prospective goals and evaluate the distance to goal locations.

As in the maze foraging tasks simulated here, one of our fMRI studies required participants to search visually for the shortest path to a goal location in novel mazes containing one (shallow maze) or two (deep maze) choice points or subgoals (Kaplan et al., 2017a). Interestingly, we observed two anterior prefrontal responses to demanding choices at the second choice point. One in rostro-dorsal medial prefrontal cortex (rd-mPFC) that was also sensitive to (deactivated by) demanding initial choices and another in lateral frontopolar cortex, which was only engaged by demanding choices at the second choice point. This suggests that, in deep mazes, these regions are engaged by belief updating during planning to identify the most promising subgoal (Kaplan et al., 2017b). Subsequent work could potentially use the active inference scheme above to fit different anterior prefrontal responses – and how they reflect robust subgoal identification. Interestingly, a recent modelling initiative showed that strong grid cell representations could lead to better calculation of subgoals, when navigating an environment (Stachenfeld et al., 2017). Although we did not measure robust entorhinal cortex signals in our experiment, our spatial planning fMRI study revealed increased hippocampal coupling with rd-mPFC when subgoals had to be identified. In the future, we hope to use the model described in this paper to elucidate the precise computational roles of the hippocampus, entorhinal cortex, and anterior prefrontal regions when formulating plans in novel environments.

A key challenge in the computational neuroscience of reinforcement learning is real world learning, in state spaces that are high-dimensional, continuous and partially observable. To meet this challenge, a recent proposal endowed reinforcement learning systems with episodic memory (Gershman and Daw, 2017). Interestingly, this approach produces equations that are formally similar to the updates in active inference (because they both involve message passing among representations of hidden or latent states in the future past). This speaks to the construct validity of both approaches, in terms of each other, and the possibility that reinforcement learning with episodic memory can be cast as active inference – and *vice versa*. Given the overlap between neuronal systems guiding spatial navigation and episodic memory (Burgess et al., 2002), an interesting future area of study will be to create agents that can navigate one environment and draw upon their previous experience when exploring another.

## Conclusion

In conclusion, we have described an active inference scheme for epistemic foraging and goal directed navigation using a minimal setup. The key contribution or insight afforded by these simulations is to show that purposeful, goal directed behaviour can be prescribed through simple prior beliefs about the outcomes that will be encountered under allowable policies. Furthermore, we have described a plausible process theory for exploratory behaviour and associated neurophysiological responses that can be tested empirically in behaving subjects. An interesting aspect of these simulations is the pursuit of long-term goals under the constraint of short-term policies. The apparent problem of failing to use appropriately distant policy horizons (i.e., deep tree searches) is easily finessed by contextualising prior beliefs such that they naturally offer attainable subgoals. Finally, we have shown that the resulting decomposition of long-term goals can operate online, even in the context of epistemic foraging. This means that goal directed behaviour and the resolution of uncertainty work hand-in-hand to underwrite predictable (minimally surprising) outcomes.

## Software note

Although the generative model – specified by the (**A, B, C,D**) matrices – changes from application to application, the belief updates in Figure 2 are generic and can be implemented using standard routines (here **spm_MDP_VB_X.m**). These routines are available as annotated Matlab code in the SPM academic software: http://www.fil.ion.ucl.ac.uk/spm/. The simulations in this paper can be reproduced (and customised) via a graphical user interface: by typing >> DEM and selecting the **Maze learning** demo.

## Acknowledgements

KJF is funded by the Wellcome Trust (Ref: 088130/Z/09/Z) and RK is funded by a Sir Henry Wellcome Postdoctoral Fellowship (Ref: 101261/Z/13/Z). We thank Mehdi Keramati for his invaluable perspective on planning and tree searches and summarising the state-of-the-art in this field. We would also like to thank our anonymous reviewers for help in describing and contextualising this work.

## Appendices

### Appendix 1 Expected free energy

variational free energy is a functional of a distribution over states, given observed outcomes. We can express this as a function of the sufficient statistics of the posterior:

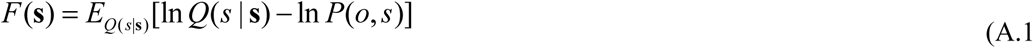

In contrast, the expected free energy is the average over (unobserved) outcomes, given some policy that determines the distribution over states. This can be expressed as a function of the policy:

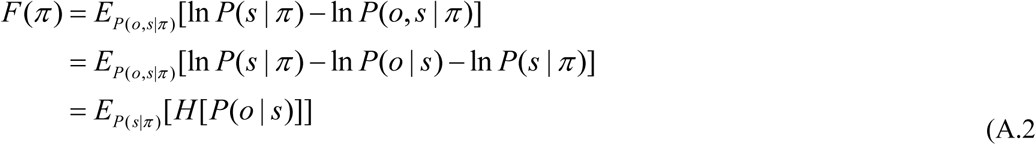

The expected free energy is therefore just the expected entropy or uncertainty about outcomes under a particular policy. Things get more interesting if we express the generative model-terms of a prior over outcomes that does not depend upon the policy

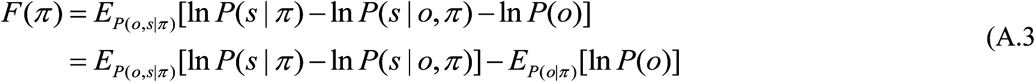

This is the form used in active inference, where all the probabilities in (A.3) are conditioned upon past observations. This enables one to replace the posterior in (A.3) with the approximate posterior that minimises variational free energy based on (observed) outcomes in the past: see (A.5).

### Appendix 2 Belief updating

Bayesian inference corresponds to minimising variational free energy, with respect to the expectations that constitute posterior beliefs. Free energy can be expressed as the (time-dependent) free energy under each policy plus the complexity incurred by posterior beliefs about (time-invariant) policies and parameters, where (with some simplifications):

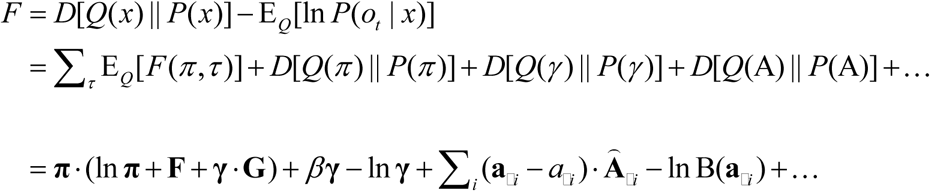

The free energy of hidden states in this expression is given by:

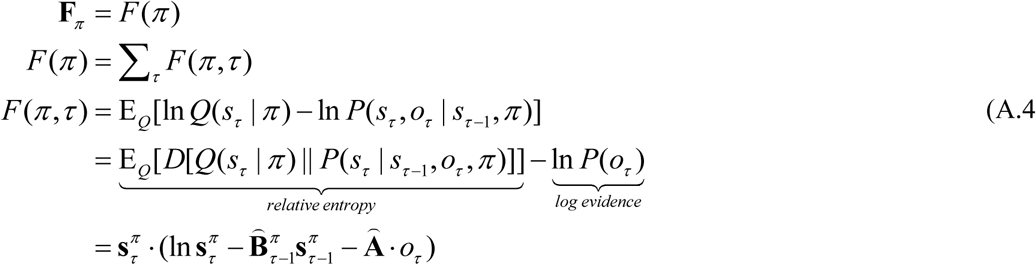

The expected free energy of any policy has a homologous form but the expectation is over both hidden states and – yet to be observed – outcomes 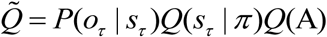:

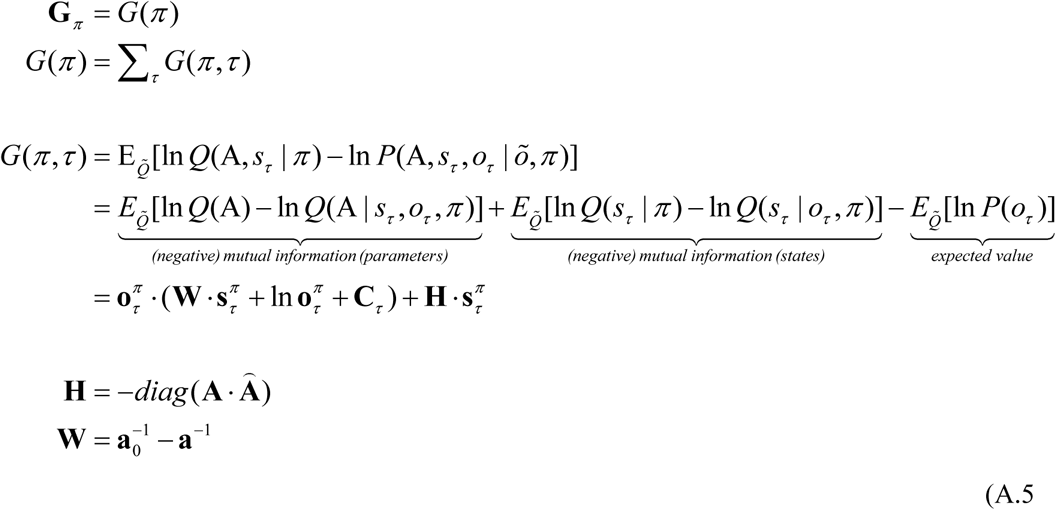

Figure 2 provides the update rules based upon minimising variational free energy via a gradient descent:

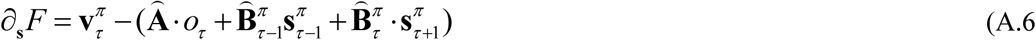

The auxiliary variables 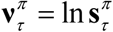 can be regarded as transmembrane potentialing a biological setting, while the resulting firing rate is a sigmoid function of depolarisation. A similar formalism can be derived for the precision (c.f., dopamine) updates. The remaining update rules are derived in a straightforward way as the solutions that minimise free energy explicitly.

